# Human body-site-stratified fungal genomic catalogs enhancing human mycobiome analysis

**DOI:** 10.64898/2026.09.20.752968

**Authors:** Sehun Ahn, Jungyeon Kim, Insuk Lee

## Abstract

Current human mycobiome profiling relies largely on marker-based methods with limited sensitivity, whereas genome-based approaches require computationally intensive alignment against broad fungal references. Here, we present MycobiomeDB, a unified fungal genome catalog, and derive body-site-specific catalogs for the gut, oral cavity, skin, and vagina through metagenome-guided stratification. Using these catalogs, we developed site-specific MycoProfilers that improved taxonomic recall and better distinguished disease-associated mycobiomes from healthy controls than existing methods.

## Main Text

Although fungi represent a relatively small fraction of the human microbiome, they can substantially influence disease-associated microbial ecosystems, making accurate mycobiome profiling clinically important^1^. Bacterial metagenomic profiling has advanced markedly through improved algorithms and comprehensive body-site-specific reference genome catalogs^2-5^. Genome-based profiling can provide greater sensitivity and taxonomic resolution than marker-based approaches, but comparable fungal resources remain limited. Comprehensive, curated fungal reference catalogs tailored to major human body sites are therefore needed to advance human mycobiome profiling.

Here, we present MycobiomeDB, a unified fungal genome catalog, and derive body-site-stratified reference catalogs for the gut, oral cavity, skin, and vagina. We compiled 16,365 fungal assemblies from NCBI GenBank and the JGI Genome Portal, retained 13,888 near-complete assemblies using EukCC^6^ (**Supplementary Fig. 1a-b**), and selected 4,094 species-representative genomes for MycobiomeDB (**Fig. 1a**). Because these genomes originate from diverse habitats, body-site assignment was based on direct metagenomic evidence from each human niche. For metagenome-guided habitat stratification, we analyzed 24,282 public whole-metagenome shotgun sequencing (WMS) samples from 125 datasets, including 9,241 gut, 2,461 oral, 9,187 skin, and 3,393 vaginal samples (**Fig. 1b, Supplementary Table 1a-d**). Kraken2^7^ was used for initial screening, followed by Bowtie2^8^-based genome-wide coverage validation. Species with ≥0.05% genome breadth and ≥1% prevalence within each body site were retained (**Supplementary Fig. 1c**), yielding 1,005 gut, 320 oral, 714 skin, and 138 vaginal species. The majority (∼75%) of fungal genomes passing the filtering criteria showed >99% completeness and <0.5% contamination (**Supplementary Fig. 1d**). We then applied conservative ecological curation using FUNGuild^9^ annotations and literature evidence, excluding taxa whose known ecology was incompatible with human association while retaining those with documented or plausible human exposure (**Supplementary Table 2**). The final body-site-specific catalogs, termed Mycobiome-HG (human gut), Mycobiome-HO (human oral), Mycobiome-HS (human skin), and Mycobiome-HV (human vagina), comprised 766, 259, 642, and 112 species, respectively (**Supplementary Table 3**).

**Figure 1.**
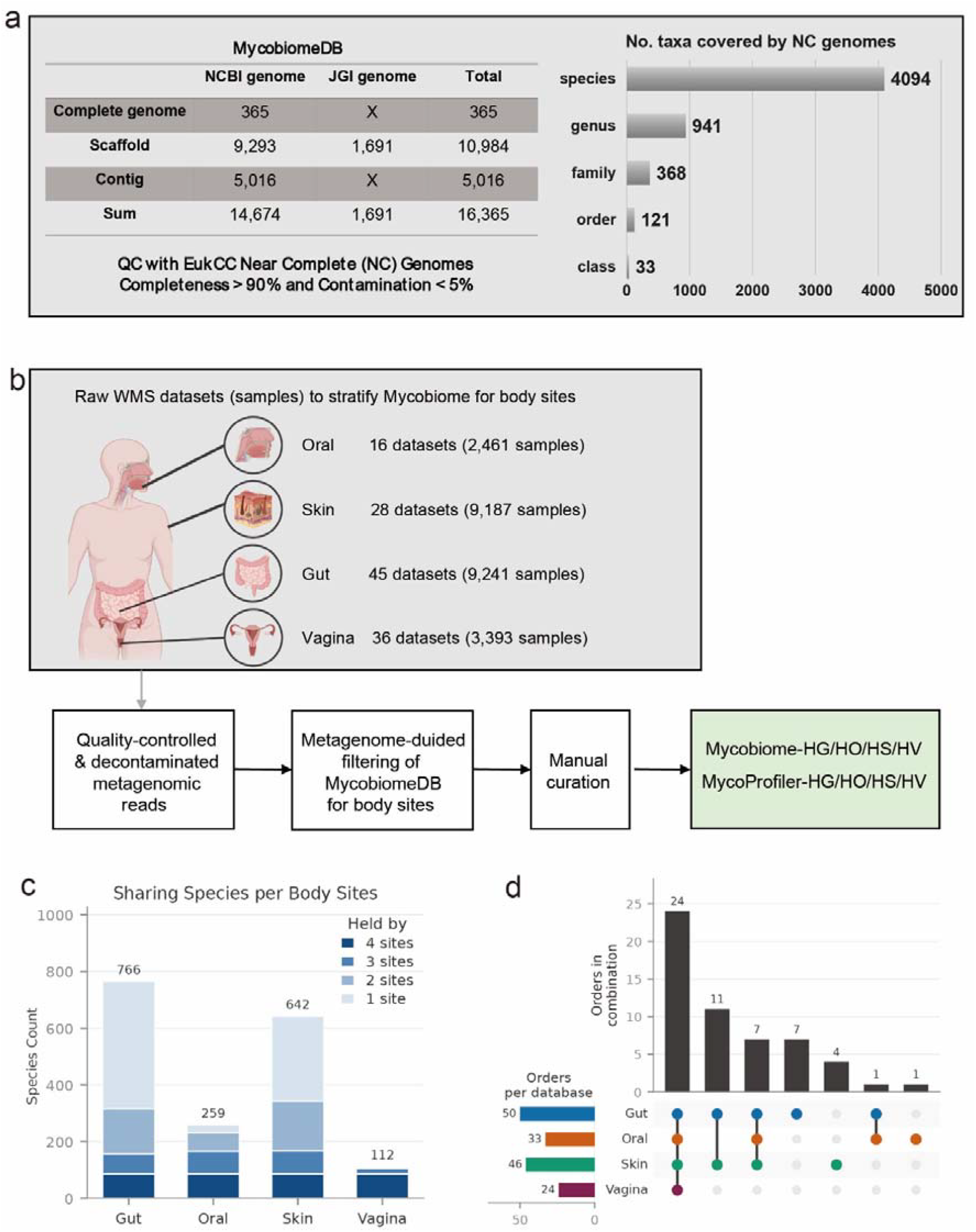
Overview of MycobiomeDB and body-site stratification. **a**, Construction of the unified fungal genome catalog, MycobiomeDB, comprising 4,094 near-complete species-representative genomes selected from 16,365 assembled fungal genomes obtained from NCBI and JGI. The bar plot shows the number of taxa at each taxonomic rank. **b**, Public whole-metagenome shotgun (WMS) datasets and workflow used to derive body-site-specific catalogs from MycobiomeDB. **c**, Species counts in the four body-site-specific catalogs, Mycobiome-HG (gut), Mycobiome-HO (oral), Mycobiome-HS (skin), and Mycobiome-HV (vagina), stratified by the number of body sites in which each species was retained. Numbers above the bars indicate the total number of species in each catalog. **d**, UpSet plot showing order-level overlap among the four body-site-specific catalogs. Horizontal bars indicate the number of orders in each catalog, and vertical bars indicate intersection sizes.

The resulting taxonomic structure strongly supported a site-aware reference design. Across the four catalogs, 1,150 distinct species were represented, of which only 86 (7%) occurred at all four sites, whereas 777 (68%) were restricted to a single catalog (**Fig. 1c**). Among the 766 gut species, 450 were gut-specific, while 299 of 642 skin species were unique to the skin catalog. In contrast, only 28 oral species were site-specific, and none of the vaginal species were unique; notably, 86 of 112 vaginal species belonged to the core shared across all four sites. These patterns indicate that the gut and skin harbor more distinct fungal communities, whereas the oral and vaginal mycobiomes show greater overlap with other body sites. Taxonomic overlap increased at higher ranks, with 24 of 55 orders shared across all four sites (**Fig. 1d**).

Site-restricted taxa were also ecologically coherent. Gut-specific groups included food- and fermentation-associated fungi, consistent with recurrent dietary passage into fecal samples^10^, whereas skin-specific groups included keratin-associated and environmentally exposed fungi, consistent with the strong topographic and environmental structuring of the skin mycobiome^11^. These patterns should not be interpreted as evidence of stable colonization; rather, the body-site catalogs in MycobiomeDB are intended to capture fungi that are empirically detectable and ecologically plausible at each site.

Next, we developed body-site-aware fungal taxonomic profilers using Mycobiome-HG/HO/HS/HV. To reduce bacterial misclassification, these profilers first conduct Kraken2 mapping of metagenomic reads against body-site-specific bacterial genome catalogs (**Methods**). Following the depletion of bacterial reads, fungal taxonomic profiling based on the Kraken2 database for each Mycobiome-HG/HO/HS/HV combined with Bracken re-estimation^12^ performed. These fungal profilers are termed MycoProfiler-HG (gut), MycoProfiler-HO (oral), MycoProfiler-HS (skin), and MycoProfiler-HV (vagina).

We compared MycoProfiler with MiCoP^13^ and EukDetect^14^, which were reported as the most accurate dedicated shotgun fungal profilers by a recent benchmark study^15^. MiCoP performs whole-genome BWA-MEM mapping followed by post-alignment evidence filtering, whereas EukDetect maps reads to a curated set of conserved eukaryotic marker genes and applies alignment-quality and multi-marker detection criteria. Because profiling performance depends on both the inference method and the underlying reference database, each profiler was evaluated using its built-in fungal reference and default parameters. We did not use simulated metagenomes as the primary benchmark because simulations generated from a common reference can assess differences in alignment or classification strategy but cannot fairly evaluate differences arising from the composition and coverage of the profilers’ native reference databases, which are a central component of their performance. As no ground truth is available for real metagenomic samples, we therefore assessed each profiler by its ability to capture disease-associated differences in mycobiome composition relative to healthy controls across 44 gut, 12 oral, 11 skin, and 5 vaginal metagenomic cohorts.

For each cohort, species-level profiles were converted to Bray–Curtis distances, and disease– control differences were assessed by permutational multivariate analysis of variance (PERMANOVA). Across body sites, MycoProfilers identified significant compositional differences between diseased and healthy samples in more cohorts than MiCoP or EukDetect (**Fig. 2a-d, e**). MycoProfilers also produced non-empty fungal profiles for a larger fraction of samples, particularly in oral and skin cohorts (**Fig. 2f**). For example, in a rheumatoid arthritis gut cohort^16^, MycoProfiler-HG distinguished diseased from healthy samples, whereas MiCoP did not, and EukDetect yielded empty profiles for most samples (**Fig. 2g**). MycoProfilers were also approximately 9-to 32-fold faster than MiCoP (Wilcoxon signed-rank *P* = 7.5 × 10□□) and had runtimes comparable to the marker-based EukDetect (**Fig. 2h; Supplementary Table 4**).

**Figure 2.**
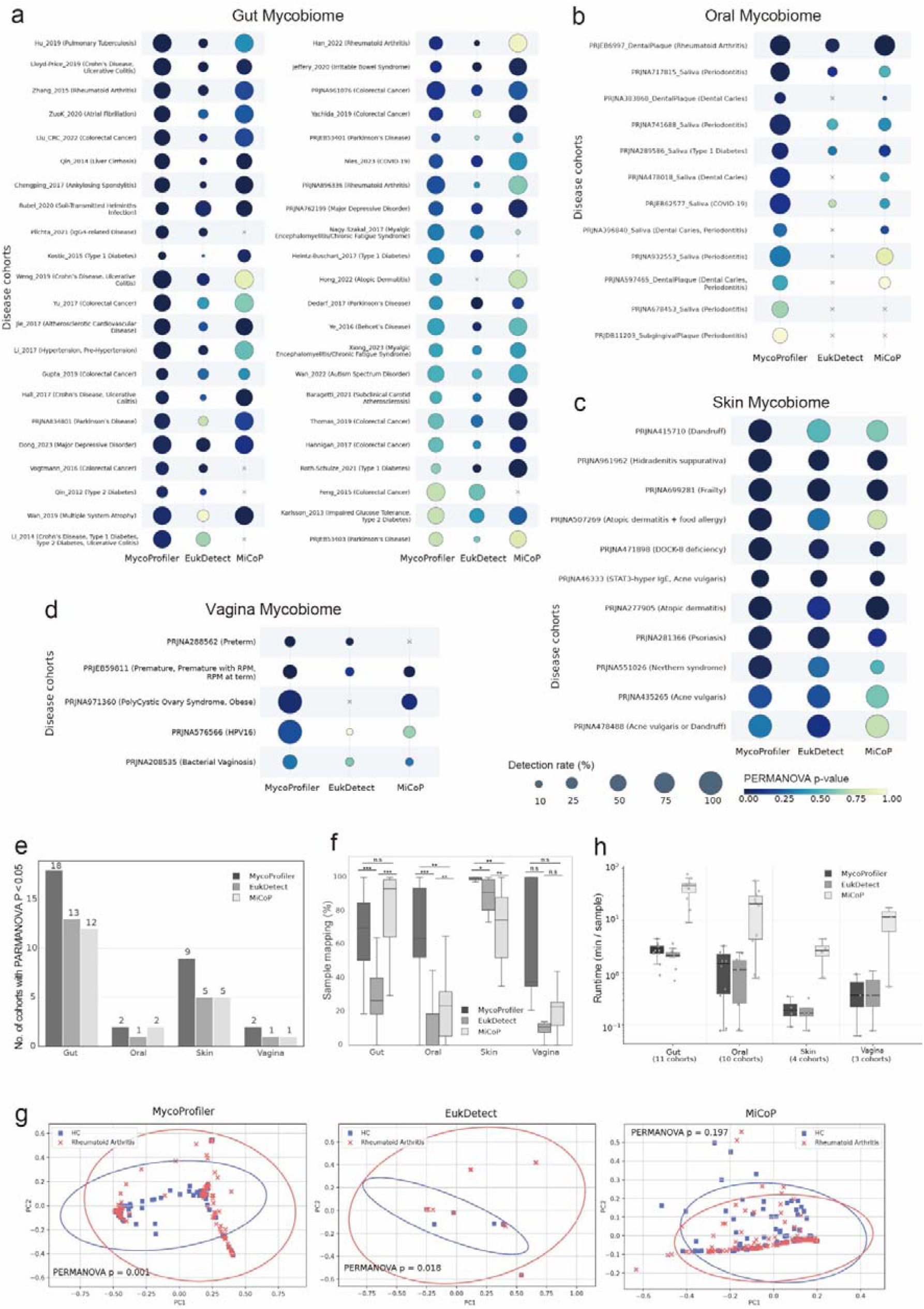
Comparison of MycoProfiler with existing fungal species profilers, EukDetect and MiCoP. **a–d**, Ability to distinguish diseased mycobiomes from healthy controls across disease-associated cohorts from the gut (a), oral cavity (b), skin (c), and vagina (d), assessed by PERMANOVA on Bray–Curtis distances. Color intensity indicates the PERMANOVA P value, and circle size represents the fraction of samples for which each profiler generated a non-empty fungal profile. **e**, Number of cohorts in which diseased and healthy mycobiomes were significantly distinguishable by each profiler (PERMANOVA P < 0.05) across body sites. **f**, Distribution across cohorts of the percentage of samples yielding a non-empty fungal profile. Differences between profilers were assessed using the Wilcoxon signed-rank test. n.s., not significant; *, P < 0.05; **, P < 0.01; ***, P < 0.001. **g**, Representative principal-coordinate analysis (PCoA) of Bray–Curtis distances for the Zhang_2015 rheumatoid arthritis cohort profiled by each method. **h**, Per-sample runtime of each profiler across body sites. Boxes indicate the interquartile range, center lines indicate medians, and whiskers extend to 1.5× the interquartile range. All tools were run on identical FASTQ inputs using 10 threads, with runtime measured using /usr/bin/time -v.

Several limitations should be considered when interpreting these results. First, the four body-site collections differed in sample size, sequencing depth, and inclusion criteria, which may have influenced the number of taxa retained in each catalog. Second, the coverage and prevalence thresholds may exclude rare but genuine fungal species from a given body site, while ecological curation is limited by incomplete annotation of fungal ecology. Third, PERMANOVA identifies differences in multivariate community structure but can be influenced by dispersion and cohort-level covariates and does not establish biological causality. Fourth, low-biomass fungal signals are particularly susceptible to reagent and laboratory contamination, and most public cohorts lacked matched negative controls^17^. Accordingly, the larger number of significant cohorts identified by MycoProfilers should be interpreted as evidence that the workflow better preserves potentially informative fungal signals, rather than as independent validation of specific disease associations.

Despite these limitations, our habitat-stratified reference strategy reduces irrelevant taxonomic search space while retaining a broad set of fungi supported by metagenomic evidence across four major human body sites. Future releases should incorporate newly sequenced isolates and fungal metagenome-assembled genomes, expand the metagenomic datasets used for habitat stratification, and undergo validation in prospective mock-community and clinical cohorts using matched negative controls and orthogonal fungal measurements.

## Methods

### Collection and taxonomic annotation of fungal genomes

As of January 17, 2024, we downloaded all fungal genomes available in NCBI GenBank, including complete, scaffold-, and contig-level assemblies (**Supplementary Table 3**). Following FunOMIC^18^, genomes with N50 <10 kb were excluded, leaving 14,674 NCBI fungal genomes. To expand genome coverage, we additionally retrieved 1,691 scaffold-level fungal genomes from the JGI Genome Portal using Advanced Search with Taxonomy = “Eukaryota,” Project Status = “Complete,” and Scientific Program = “Fungal.” Genomes lacking NCBI taxonomy assignments were excluded. NCBI taxonomy dump files were then used to annotate all retained genomes across the standard taxonomic hierarchy: kingdom, phylum, class, order, family, genus, and species.

### Genome quality assessment and species-level dereplication

To retain near-complete genomes, we assessed genome quality using EukCC^6^ with the --DNA and --clade fungi options (**Supplementary Fig. 1a,b**). Genomes with ≥90% completeness and ≤5% contamination were retained.

To reduce database redundancy, we selected one representative genome per species. For species represented by multiple genomes, the genome with the highest genome intactness score (*S*)^5^, *S* = *Completeness* − 5 × *Contamination* + 0.5 × log_10_ (*N50*), was retained; ties were resolved by selecting the genome with the longest assembly length. This procedure yielded a pre-filtered database of 4,094 high-quality species-representative fungal genomes.

### Collection and curation of body-site-aware metagenomic datasets

Publicly available whole-metagenome shotgun sequencing (WMS) datasets from the human gut, oral cavity, skin, and vagina were collected from the NCBI Sequence Read Archive (SRA) and European Nucleotide Archive (ENA). Datasets were identified using two complementary strategies: literature searches in PubMed and Google Scholar for published human WMS studies, and direct searches of the SRA and ENA to capture additional published and unpublished datasets. Literature searches combined body-site, disease phenotype, and sequencing-related keywords. For ENA Advanced Search, “Raw reads” was selected as the datatype, with library layout = PAIRED, library strategy = WGS, and library source = GENOMIC.

Because dataset availability and sequencing characteristics differed across body sites, site-specific inclusion criteria were applied. Gut datasets were restricted to paired-end samples with ≥3 Gbp sequencing depth, whereas all paired-end oral samples were included regardless of depth. For skin and vaginal datasets, all available WMS samples were retained without additional depth filtering.

In total, 24,282 WMS samples from 125 datasets were collected, comprising 9,241 gut samples from 45 datasets, 2,461 oral samples from 16 datasets, 9,187 skin samples from 28 datasets, and 3,393 vaginal samples from 36 datasets (**Supplementary Table 1a-d**).

### WMS read preprocessing and host decontamination

WMS reads were preprocessed by adaptor trimming and low-quality read removal using Trimmomatic (v.0.39)^19^ with the appropriate adaptor sequences. Host-derived reads were then removed by aligning the quality-filtered reads to the human reference genomes GRCh38.p13^20^ and T2T^21^, using Bowtie2 (v.2.5.4)^8^. Reads mapping to either human reference genome were excluded from downstream analyses.

### Metagenome-guided stratification of body-site-specific fungal genome catalogs

Because MycobiomeDB contains fungal genomes recovered from diverse habitats, we defined body-site-specific catalogs based on direct evidence of detection in metagenomic reads from each human body site, assuming that genomes reproducibly supported by reads from a given site are likely to represent fungi present in that ecological niche. We therefore applied a two-stage filtering strategy to 9,241 gut, 2,461 oral, 9,187 skin, and 3,393 vaginal metagenomes: Kraken2 (v2.1.6)^7^ was first used for sensitive species detection at a confidence threshold of 0.2, after which all species detected within each cohort were pooled to construct a cohort-specific Bowtie2 (v2.5.4)^8^ reference for genome-wide read alignment and subsequent validation of species presence.

Genome breadth of coverage for each fungal species was quantified per sample using samtools^22^ coverage. To define empirical retention criteria, we examined the number of retained species across increasing breadth-of-coverage thresholds from 0% to 1% in 0.01% increments and prevalence thresholds from 0% to 5% in 0.1% increments. Based on these distributions, species were retained when they showed ≥0.05% genome breadth and ≥1% prevalence within the corresponding body-site collection (**Supplementary Fig. 1c**). This hybrid strategy reduced the search space and limited spurious marker-based assignments while preserving species supported by recurrent genome-wide read evidence.

### Manual ecological curation of body-site-associated fungal taxa

Following computational filtering, we performed conservative manual curation to remove fungal taxa with ecologies clearly incompatible with human body sites while preserving potentially relevant species. Curation was conducted primarily at the family level because ecological information was more consistently available at this rank and related taxa often share similar trophic modes. Nutritional modes were assigned using FUNGuild^9^ and evaluated for compatibility with human-associated environments.

Among pathotrophs, taxa primarily associated with insect infection were excluded. Among symbiotrophs, groups restricted to wood, soil, or lichen-associated niches were removed. For saprotrophs, edible mushroom taxa were retained, whereas inedible or toxic mushroom taxa were excluded (**Supplementary Table 2**). To avoid excessive filtering, taxa were retained if they had any documented detection in humans, were reported in studies involving human samples, or had insufficient ecological information for confident exclusion. This conservative curation was intended to improve biological plausibility without sacrificing sensitivity, recognizing that taxa not genuinely present in the human microbiome would receive little or no read support in downstream profiling.

### Development of body-site-aware fungal species profilers

Body-site-aware fungal profilers were developed using the corresponding Mycobiome-HG/HO/HS/HV reference catalogs. To reduce bacterial misclassification, metagenomic reads were first depleted against body-site-specific human bacterial genome catalogs using Kraken2^7^: Human Reference Gut Microbiome (HRGM)^5^ for gut, Human Reference Oral Microbiome (HROM)^4^ for oral, Skin Microbial Genome Collection (SMGC)^2^ for skin and Vaginal Microbial Genome Collection (VMGC)^3^ for vaginal samples. The remaining reads were then classified against the corresponding Mycobiome-HG/HO/HS/HV catalog with Kraken2 at a confidence threshold of 0.2, and species abundances were re-estimated using Bracken^12^. The resulting profilers were termed MycoProfiler-HG, MycoProfiler-HO, MycoProfiler-HS, and MycoProfiler-HV for the human gut, oral cavity, skin, and vagina, respectively.

### Benchmarking of fungal taxonomic profilers

We benchmarked MycoProfilers against two existing fungal profilers, MiCoP^13^ and EukDetect^14^, using real metagenomic samples. Because the three tools rely on different fungal reference databases, direct benchmarking with a common simulated community was not considered appropriate. MiCoP and EukDetect were run with default parameters against their built-in reference databases.

For MycoProfilers, Bracken^12^ read length was matched to each cohort. Because Bracken requires a k-mer distribution generated for a specific read length, we precomputed distributions for 64, 70, 77, 90, 100, 125, 136, 150, 200, and 250 bp. The representative read length of each cohort was determined from its FASTQ files, and the closest precomputed distribution was used for abundance estimation.

Species-level relative abundance profiles generated by each tool were converted to Bray– Curtis distance matrices using the beta_diversity function in the skbio.diversity module of the scikit-bio. Differences in community composition between groups were tested by permutational multivariate analysis of variance (PERMANOVA) using skbio.stats.distance.permanova.

Computational performance was evaluated by executing all three tools on the same quality-controlled FASTQ files using 10 threads, matching the thread settings used for the archived MiCoP (--threads 10) and EukDetect (--cores 10) runs. Wall-clock time was measured separately for each cohort by wrapping the complete cohort-level profiling workflow with /usr/bin/time -v and extracting the value reported in the “Elapsed (wall clock) time” field. The same timing instrumentation and parsing procedure was applied consistently to MycoProfilers, MiCoP, and EukDetect. Because runtime was measured at the cohort level, the total wall-clock time for each cohort was divided by the number of samples in that cohort to obtain a per-sample runtime for cross-cohort comparisons. Peak resident memory usage was obtained from the “Maximum resident set size” field of the same /usr/bin/time -v report.

## Data Availability

The body-site-specific catalogs Mycobiome-HG, Mycobiome-HO, Mycobiome-HS, and Mycobiome-HV are publicly available through Zenodo at https://doi.org/10.5281/zenodo.21449077. Accession information for the publicly available fungal genomes and whole-metagenome shotgun datasets used in this study is provided in the Supplementary Tables.

## Code Availability

The MycoProfiler pipelines, together with documentation and usage instructions, are available at https://github.com/netbiolab/MycobiomeDB.

## Acknowledgments

This research was supported by the National Research Foundation of Korea (NRF), funded by the Ministry of Science and ICT (MSIT) (RS-2026-25549861, RS-2025-18362970 to I.L.).

## Author contributions

S.A. and I.L. conceived the study. S.A. performed data analysis and catalog development. J.K. assisted genomic data compilation and analysis. I.L. supervised the project. S.A. and I.L. wrote the manuscript.

---

**Supplementary Figure 1.**
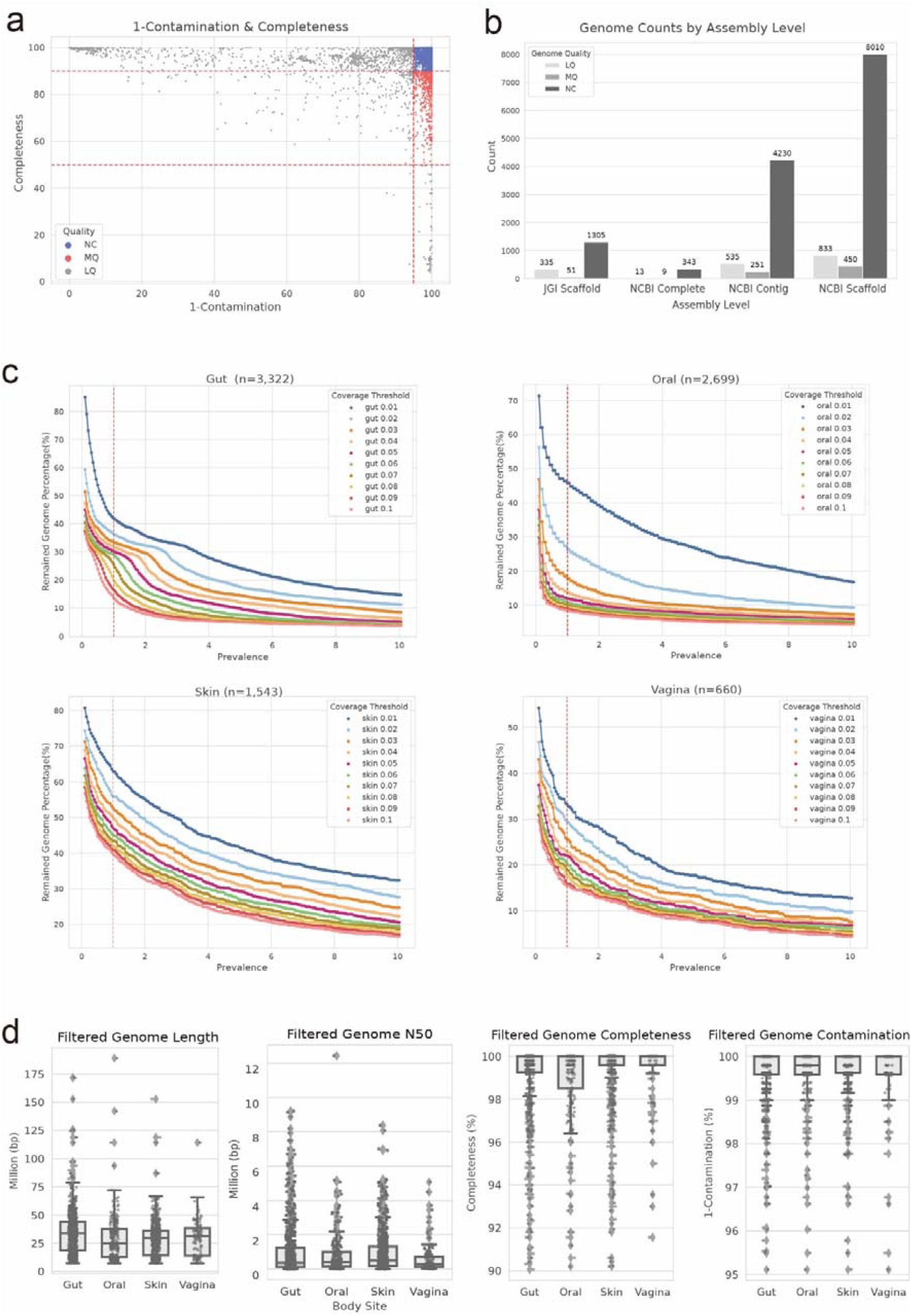
Details of metagenome-guided stratification for body-site-specific fungal genome catalogs. **a**, EukCC estimates of completeness and contamination for collected fungal assemblies. Dashed lines indicate quality thresholds: near-complete (NC), completeness ≥90% and contamination ≤5%; medium-quality (MQ), completeness ≥50% and contamination <10% but not meeting NC criteria; and low-quality (LQ), assemblies not meeting the MQ criteria. Colors denote NC, MQ, and LQ assemblies. **b**, Numbers of NC, MQ, and LQ genomes stratified by source and assembly level. **c**, Percentage of candidate genomes retained across prevalence thresholds from 0 to 10% and breadth-of-coverage thresholds from 0.01% to 0.1% in gut, oral, skin, and vaginal datasets. The vertical dashed line indicates the 1% prevalence threshold used for catalog construction. **d**, Distributions of genome length, N50, completeness, and 1 − contamination for genomes included in the four body-site-specific catalogs.

